# Study of Plasma Adiponectin Levels in Patients with Metabolic Syndrome and Therapeutic Indication in Bangladesh

**DOI:** 10.1101/2024.10.15.618480

**Authors:** Salina Shaheen Parul, Reaz Ahmmed, Md. Taohid Hasan, Ariful Islam, Md. Motiar Rahman, M. Manirujjaman, Md. Wasim Bari, Md. Sakil Ahmed, Md. Sohel Hasan, Mohammad Amirul Islam

**Author notes:** **Authors email addresses** (Salina Shaheen Parul); (Reaz Ahmmed); (Md. Taohid Hasan); (Ariful Islam) (M.Manirujjaman), (Motiar Rahman), (Md. Wasim Bari); (Md. Shakil Ahmed), (Md. Sohel Hasan). **Corresponding Author:**, Department of Biochemistry and Molecular Biology, University of Rajshahi, Rajshahi-6205, Bangladesh.

## Abstract

**Background:** The metabolic syndrome (Met-S) is a cluster of some interrelated common clinical disorders, including central obesity, dyslipidemia, hypertension, and glucose intolerance. Central obesity, accompanied by insulin resistance, is a key factor for the development of metabolic syndrome. Adiponectin is an adipose-specific plasma protein, secreted from adipocyte with anti-atherogenic and insulin-sensitizing activities.

**Purpose:** This study aimed to investigate the relationship of plasma adiponectin levels with metabolic syndrome, related disorders and its drug repurposing through in silico approach.

**Materials and Methods:** For this study, 269 individuals were recruited with written consent. The participants were selected based on their full medical history, clinical examination, and laboratory reports. Anthropometric measurements as well as blood pressure was measured before sample collection. Fasting blood samples were collected for the estimation of lipid profile, blood glucose, and serum adiponectin levels.

**Results:** Our results show that the adiponectin levels in the subjects with Met-S were significantly lower than those of without Met-S (p <0.0001). Among the metabolic syndrome risk factors, adiponectin levels were associated with hypertriglyceridemia and reduced HDL-cholesterol (p<0.0001). Three drugs (Saquinavir, Candesartan and Glimepiride) were suggested to control the plasma adiponectin level in the subjects with Met-S.

**Conclusions:** Since the plasma adiponectin levels are significantly lower in patients with Met-S, it might be used as diagnostic & prognostic marker for Met-S disorder and adiponectin targeted drugs might be minimize the Met-S of the subjects.

## 1. Introduction

The Met-S is a cluster of some physiological and anthropometric disorders and considered as notable risk factors of type II diabetes and cardiovascular diseases [1–3]. It has been reported that around 20-25% of the world’s population suffers from some types of Met-S [4]. The Third National Health and Nutrition Examination Survey revealed that greater than 20% of the US adults suffered from Met-S only between 1988 and 1994 [5,6]. Another statistics of NHANES (2003-2006) demonstrated that the incidence of Met-S was around 35% and 33% in men and women, respectively [7]. A population-based study revealed that about 19.5% rural Bangladeshi adults are sufferers of Met-S [8].

As a result of obesity, caused by modern lifestyle changes, the number of instances with Met-S has increased considerably in recent years. The patients with Met-S are more likely to have a heart attack, stroke or even death compared to the healthy individuals without Met-S[2,9,10]. However, the etiological factors responsible for the development of Met-S is still unknown. Therefore, Met-S has become one of the crucial concerns in current public health programs [11–13]. Recent research has shown that adipose tissue generates a variety of bioactive molecules known as adipocytokines, and the dysregulation of these molecules in visceral or abdominal obesity may stimulate to the progression of Met-S [14].

Adiponectin, a type of adipocytokines, is an adipose-specific protein that often exists in multimeric forms (12 to 18-mers) with varying molecular weight [15]. It has two ubiquitously expressed receptors such as, AdipoR1 and AdipoR2. AdipoR1, mostly expressed in skeletal muscle, can activate AMPK to promote lipid oxidation. On the other hand, AdipoR2 is highly expressed in liver where it enhances insulin sensitivity, and regulates glucose homeostasis [3]. Despite the fact that adiponectin is an important molecule in cells, it has been demonstrated to be underregulated as a result of a number of risk factors, including excess body mass and obesity, as well as the SNP176 genetic factor [16–19].

The deficiency of adiponectin levels appears to be a key factor in the development of insulin resistance, type 2 diabetes, dyslipidemia, and other metabolic diseases. Moreover, reduced adiponectin levels are directly related to the development of atherosclerosis [15,20–22]. For a better understanding of the relationship of Met-S with diabetes and cardiovascular disease (CVD), this protein may be considered as a promising adipokine [23]. Therefore, it is required to adiponectin-guided drug agents who are suffering from metabolic syndrome. Although adiponectin is associated with a variety of disorders, the real scenario of this protein in Bangladeshi people with metabolic disorders has not been yet elucidated. This study was designed to investigate the link between Met-S and adiponectin in the Southern Part of Bangladesh. Schematically the overview is given **Fig.1**

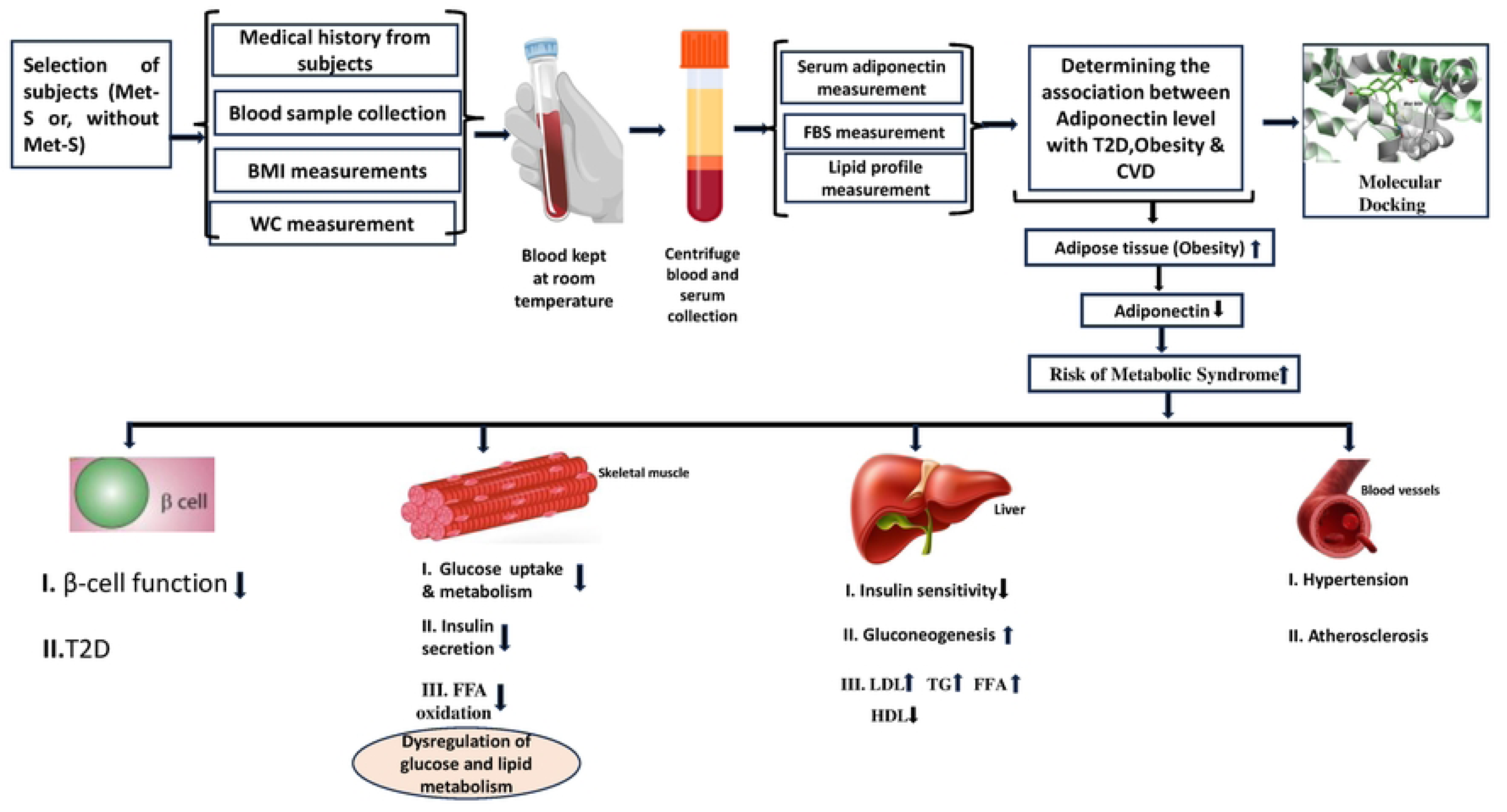

## 2. Materials and methods

### 2.1. Study Subjects

For this study 269 individuals of 30∼65 years old were recruited from the department of Medicine of Khulna Medical College Hospital, Khulna, Bangladesh. Subjects with hyper-and hypothyroidism, cardiovascular disease, acute infectious disease, chronic kidney failure, cancer, pregnancy or nursing mother, Cushing’s syndrome and any types of disability within six months were excluded from this study. Before inclusion, an informed written consent was taken from all of the subjects. All individuals with Met-S were grouped according to the modified National Cholesterol Education Program criteria [24], and were considered as Met-S subjects if at least two of the following abnormalities were present: (a) waist circumference (>90 cm for males; >80 cm for females), (b) systolic blood pressure (≥130mmHg) or diastolic pressure (≥85 mmHg), (c) triglyceride (≥150 mg/dl), (d) high-density lipoprotein cholesterol (<40 mg/dl in males; <50 mg/dl in females, and (e) fasting blood sugar (≥108 mg/dl).

### 2.2. Anthropometric Measurement

BMI was measured by dividing an individual’s weight by the square of their height in m^2^ (kg/m^2^). Waist circumference was calculated at the midpoint between the iliac crest and the lowest rib cage. A sphygmomanometer was used to measure blood pressure of the individuals in a sitting position after two minutes of rest.

### 2.3. Biochemical Analysis

For biochemical analyses, 5 ml of blood was drawn into prelabelled test tubes and was subsequently left at room temperature to coagulate the blood samples properly. The samples were centrifuged for 2 min at 3500 rpm. Serum was used to assess fasting blood sugar and lipid profiles using automatic bioanalyzer (Vitalab, Netherlands) with commercially available kits (Randox, UK). Serum adiponectin levels were measured using the Human Adiponectin Enzyme-Linked Immunosorbent Assay (ELISA) kit (Chemicon, USA).

### 2.4. Drug repurposing

To explore potential repurposable drug molecules, we performed molecular docking by using AutoDock Vina [25], drug-Likeness using SCFBio [26], ADMET using SwissADME [27], amdetSAR [28], pkCSM [29] and Protein–Ligand Interaction Profiler (PLIP) [30]. Detail descriptions are given in supplementary **Method S1**.

### 2.5. Statistical Analysis

SPSS (Statistical Package for Social Science, Chicago, IL, USA) software version 21.0 was used for statistical analysis. The value of each test was calculated as mean ± SD. The serum adiponectin levels of two groups, with or without Met-S groups, were compared using student’s t-test. Assumption of normality were assessed by visual observation of histogram with QQ plot. The Pearson correlation was employed to check whether serum adiponectin was correlated with other parameters, such as age, BMI and Met-S risk factors or not.

Univariable and multivariate linear regression were applied to determine the association among the risk factors of Met-S and serum adiponectin. In the multivariate linear regression model backward elimination procedure was used to achieve the final model. Beta coefficient (β) with standard errors (SE) and coefficient of determination (R^2^) were presented. P-value <0.05 was considered statistically significant. All the graphical illustrations were generated using GraphPad prism software.

## 3. Results

### 3.1 Clinical characteristics

**Table 1** exhibits the clinical characteristics of the subjects used in the study. The mean age of subjects with Met-S (group 1) and without Met-S (group 2) was 49.54±9.22 years and 48.05±7.54 years, respectively. The mean BMI of individual with Met-S was 25.22±2.70 46 (kg/m^2^) whereas this value for individual without Met-S was 22.85±2.22 (kg/m^2^). Waist circumference (WC) of group 1 (96.55±9.38 cm) was around 14 cm higher compared to group 2 (82.42±4.56 cm). The mean ± SD of systolic blood pressure with Met-S and without Met-S was 129.90±14.40 mmHg and 118.50±12.09 mmHg, respectively. On the other hand, diastolic pressure for individual with Met-S was 84.58±9.84 mmHg and without Met-S was 77.98±7.63 mmHg. The results indicate that the values of BMI, waist circumference, systolic and diastolic blood pressure were significantly higher than that of individuals without Met-S.

**Table 1.** Clinical and biochemical characteristics of study subjects.

| Parameters | Subjects with Met-S<br>(Group 1), n=146 | Subjects without Met-S<br>(Group 2), n=123 | t-value | P-value |
| --- | --- | --- | --- | --- |
| Age (year) | 49.54 ± 9.22 | 48.05 ± 7.54 | 1.44 | 0.1522 |
| BMI (Kg/m<sup>2</sup>) | 25.22 ± 2.70 | 22.85 ± 2.22 | 7.75 | <0.0001 |
| WC (cm) | 96.55 ± 9.38 | 82.42 ± 4.56 | 15.25 | <0.0001 |
| SBP (mmHg) | 129.90 ± 14.40 | 118.50 ± 12.09 | 6.98 | <0.0001 |
| DBP (mmHg) | 84.58 ± 9.84 | 77.98 ± 7.63 | 6.05 | <0.0001 |
| FBS (mg/dl) | 148.20 ± 62.38 | 96.10 ± 7.97 | 9.20 | <0.0001 |
| TG (mg/dl) | 277.51 ± 13.10 | 140.60 ± 20.34 | 13.24 | <0.0001 |
| HDL (mg/dl) | 33.07 ± 5.72 | 41.13 ± 4.31 | -12.85 | <0.0001 |
| TC (mg/dl) | 197.40 ± 33.26 | 171.2 ± 19.26 | 7.73 | <0.0001 |
| LDL (mg/dl) | 108.60 ± 26.93 | 99.76 ± 20.52 | 2.99 | 0.0031 |

*\*p-value < 0.05 was considered significant*

### 3.2. Biochemical characteristics

Adiponectin levels in the individuals with Met-S were lower (3.79±1.96 µg/ml) when compared to the individuals without Met-S (5.91+2.02 µg/ml) and the difference of adiponectin levels was significant between these two groups (p < 0.0001; Figure 1). The mean ± SD of Fasting Blood Sugar (FBS) (mg/dl) in participants with Met-S was 148.20±62.38 mg/dL while this value was 96.10±7.97 mg/dl for individual without Met-S. The difference of FBS concentration of these two groups was highly significant (P-value < 0.0001). The mean ± SD of triglycerides (TG) and total cholesterol (TC) levels were significantly higher in group I subjects (TG: 277.51±13.10 mg/dl; TC: 197.40±33.26 mg/dl) than group 2 (TG: 140.60±20.34 mg/dl; TC: 171.2±19.26 mg/dL). However, the high-density lipoprotein (HDL) was much lower in participants of group 1 (33.07±5.72 mg/dl) compared to subjects of group 2 (41.13±4.31 mg/dl). **Table 1** shows that that the differences of LDL levels between these were not statistically significant.

**Figure 2.**
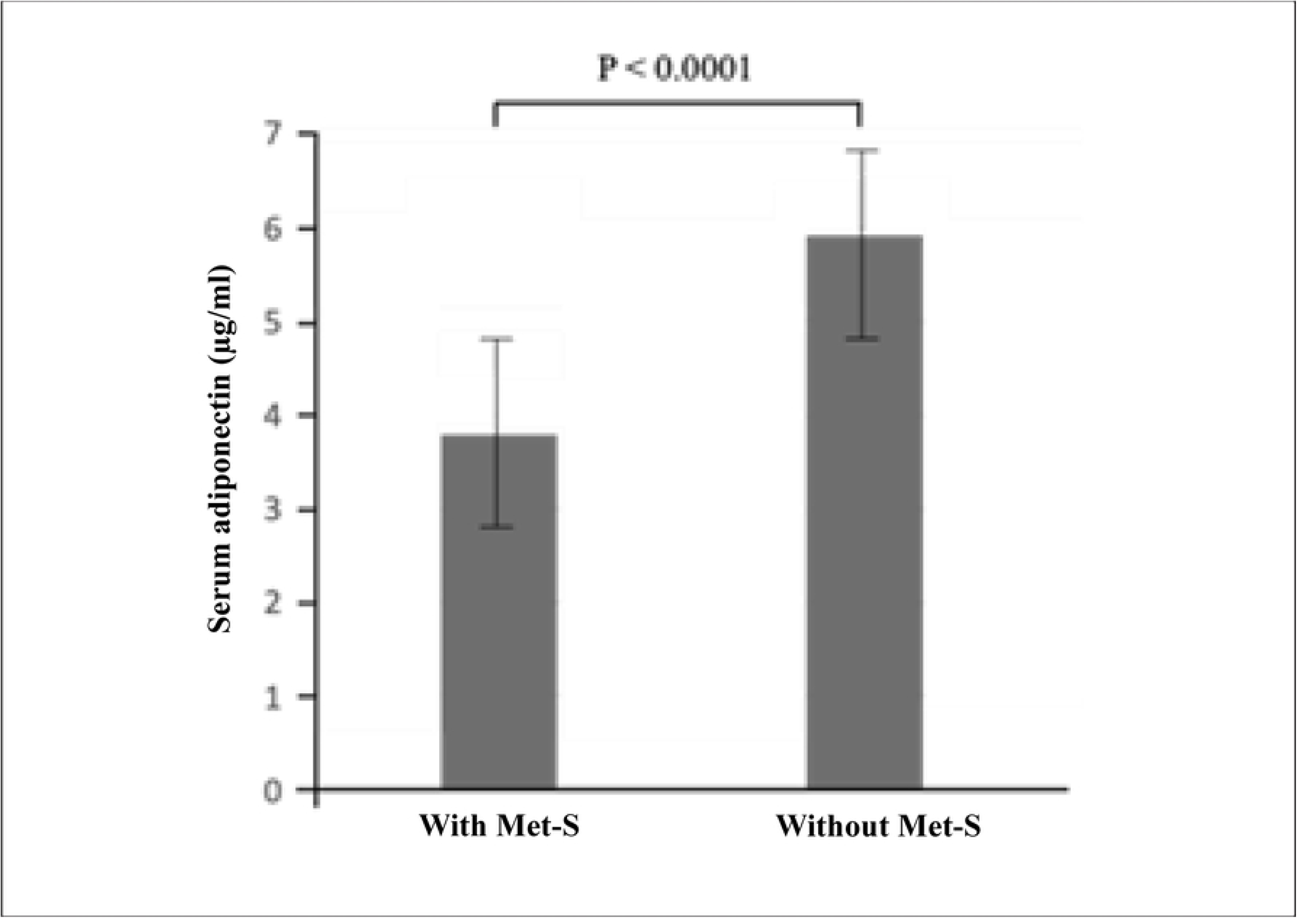
The serum levels of adiponectin levels in subjects with or without Met-S (CI 95%).

### 3.3. The association of adiponectin with metabolic risk factors

To investigate the effects of metabolic risk factors on serum adiponectin levels, unpaired students t-test was conducted. Our study shows that adiponectin levels was significantly lower in subjects with metabolic syndrome and other risk factors including obesity, elevated triglycerides, cholesterol, and hypertension (Figure 2). Subjects with central obesity, elevated triglyceride, elevated total cholesterol, and hypertension had a significantly reduced level of serum adiponectin level than normal participants (p <0.0001). Similarly, subjects with reduced High-Density Lipoprotein (HDL) levels showed significantly (p <0.0001) lower concentration (4.341 µg/ml) of serum adiponectin as compared to subjects with higher HDL (5.85 µg/ml).

**Figure 3.**
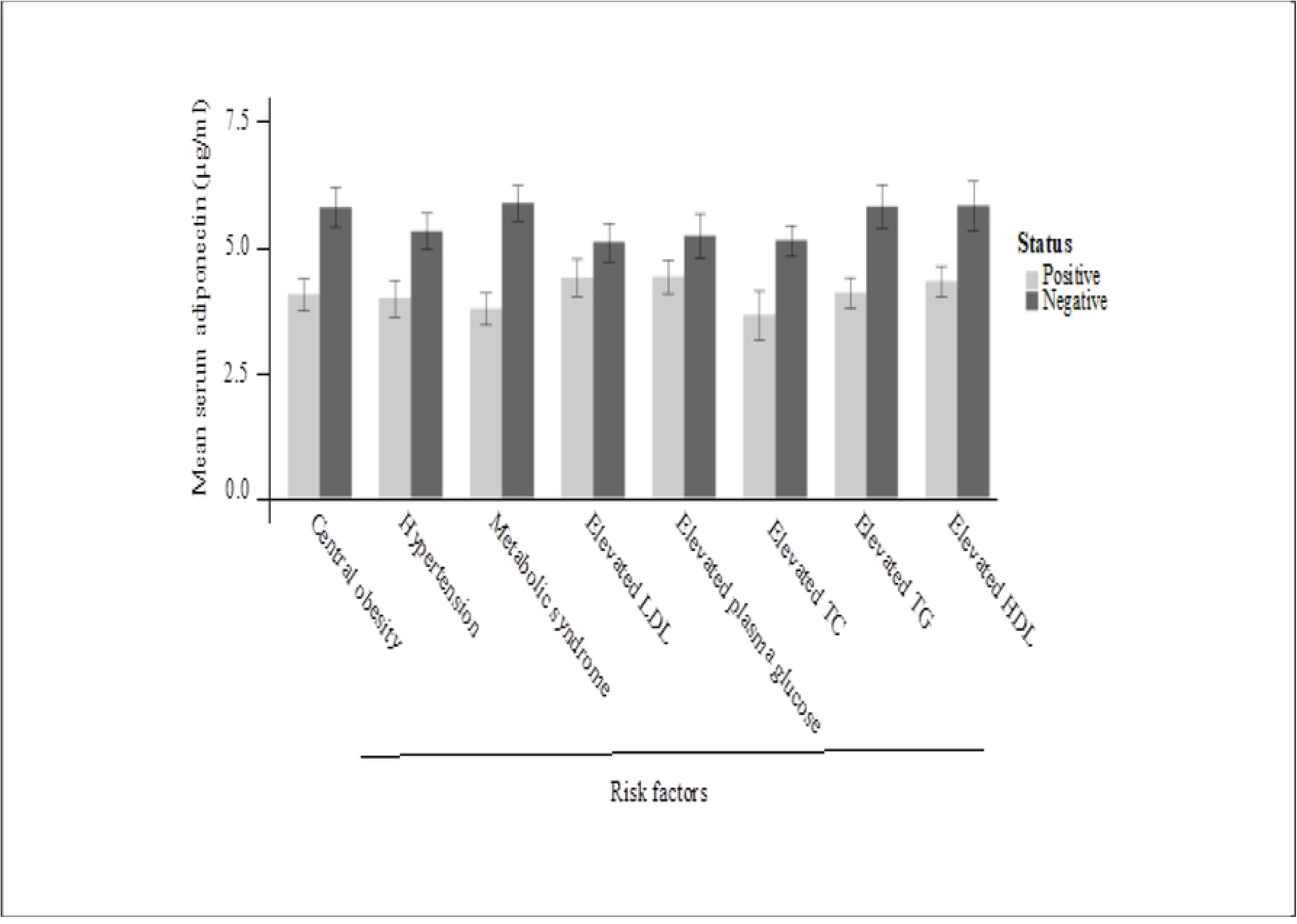
The mean of serum adiponectin concentration based on metabolic syndrome risk factors (CI 95%).

### 3.4. Correlation of adiponectin with risk factors in individual group

We next calculated the correlation coefficients (r) of serum adiponectin with clinical and biochemical parameters among group 1 and group 2 subjects separately. Our analyses showed that adiponectin has insignificant negative correlation with age, BMI and WC in group 1 subjects. However, serum adiponectin had significant negative correlation with diastolic blood pressure (r = – 0.237, p = 0.003), TG (r = –0.323, p = <0.0001) and TC (r = –0.175, p = 0.0341), respectively. Adiponectin showed insignificant negative correlation with systolic blood pressure (r = –0.1573, p = 0.0578) and LDL (r = – 0.01313, p = 0.875). A significant negative correlation was observed between adiponectin and DBP. On the other hand, a significant positive correlation was shown between adiponectin and HDL (r = 0.1764, p = 0.033) (**Table 2**).

**Figure 4.**
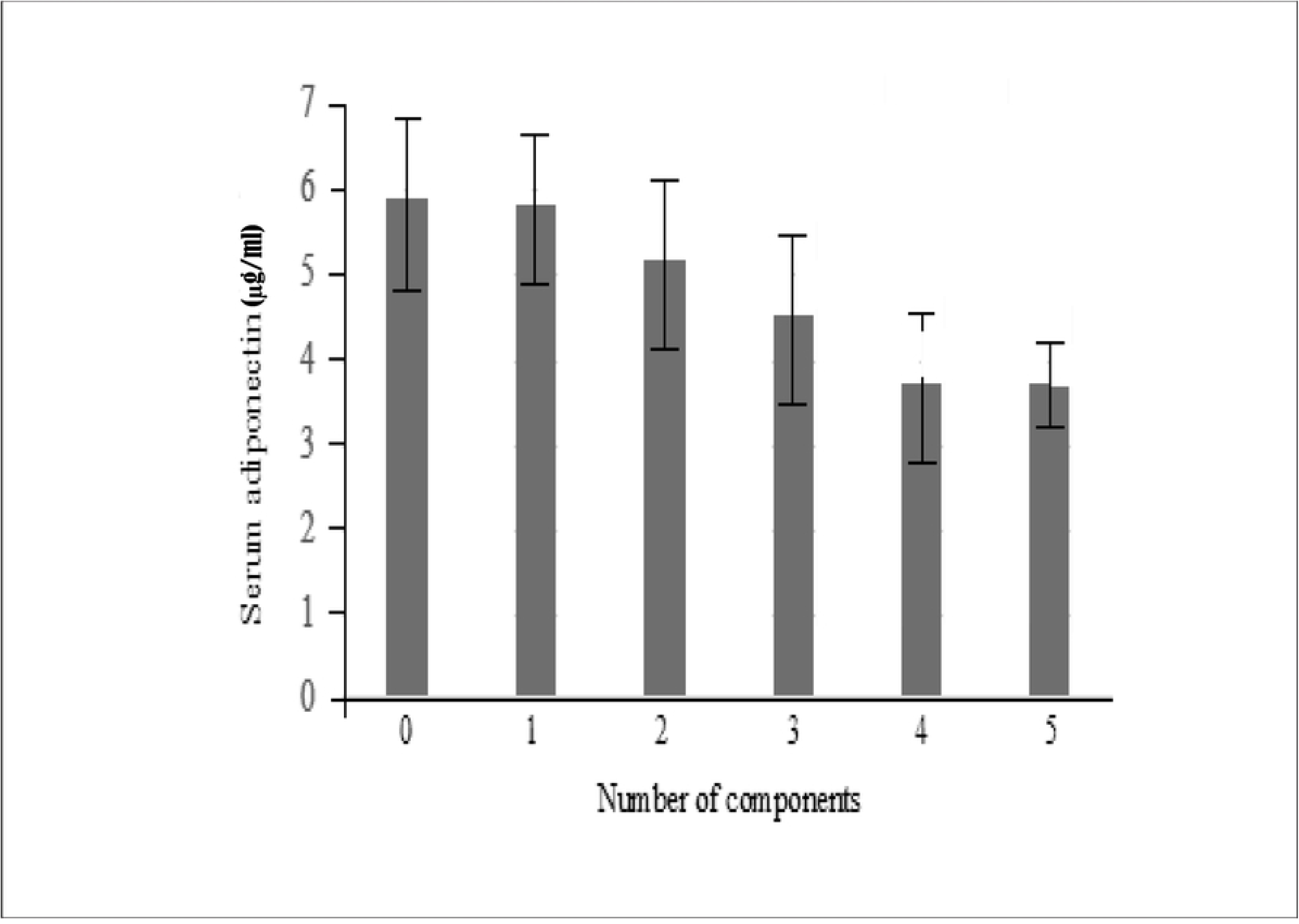
The serum adiponectin concentration with the number of Met-S components (CI 95%.

**Table 2.**
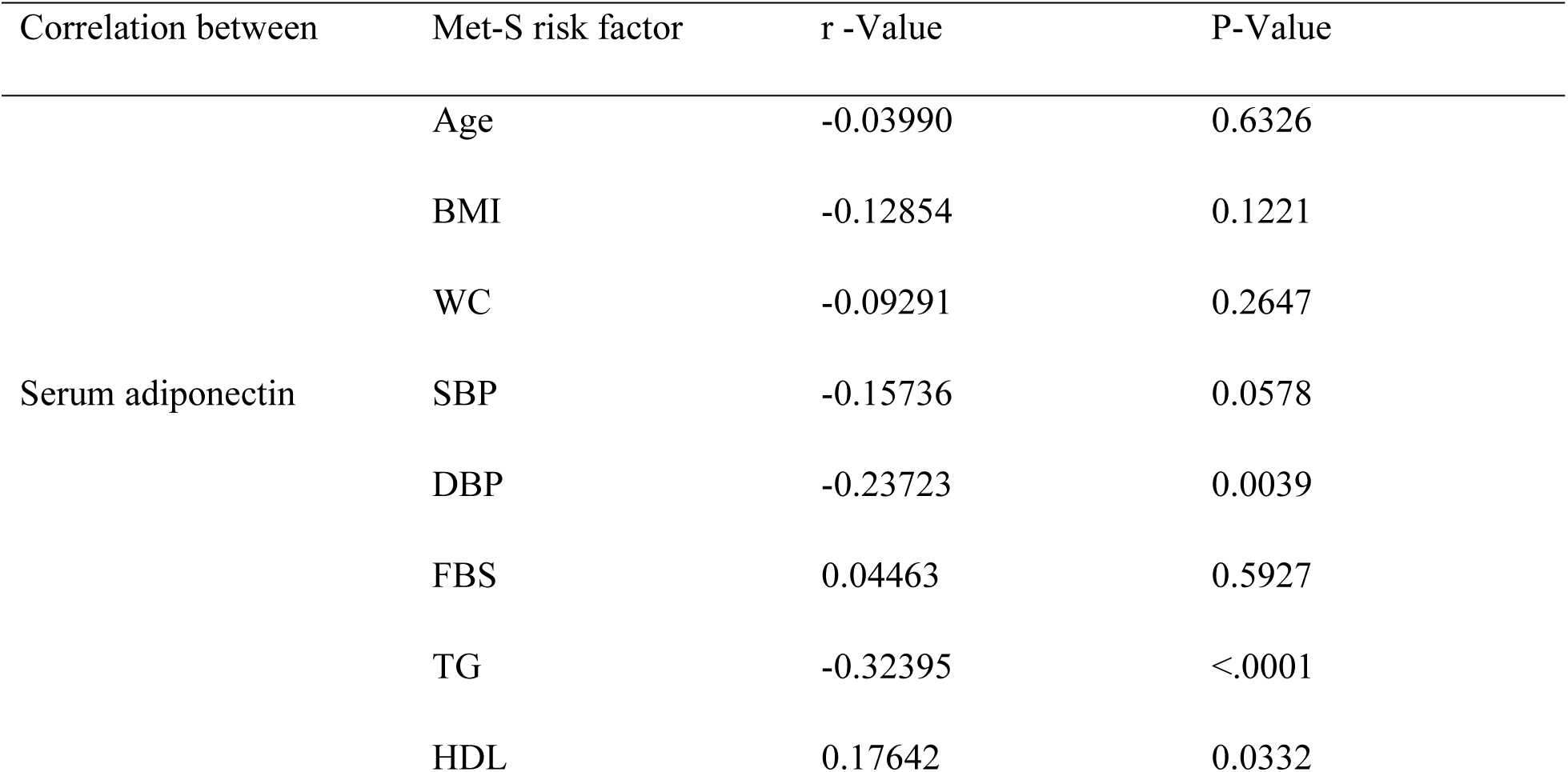
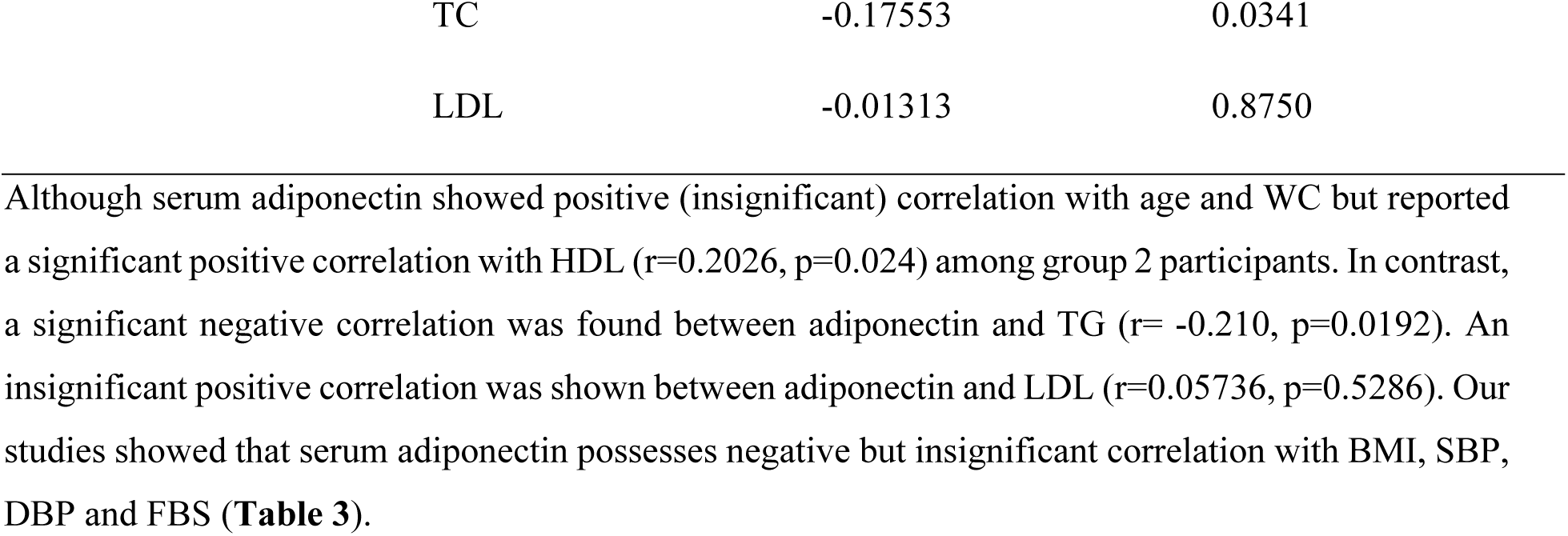
Correlation of S. adiponectin with Age, BMI and Met-S risk factor in Group –1.

**Table 3.** Correlation of serum adiponectin with Age, BMI and Met-S risk factor in Group –2.

| Correlation between | Met-S risk factor | r-Value | P- Value |
| --- | --- | --- | --- |
| Serum adiponectin | Age | -0.03990 | 0.6326 |
|  | BMI | -0.12854 | 0.1221 |
|  | WC | -0.09291 | 0.2647 |
|  | SBP | -0.15736 | 0.0578 |
|  | DBP | -0.23723 | 0.0039 |
|  | FBS | 0.04463 | 0.5927 |
|  | TG | -0.32395 | <.0001 |
|  | HDL | 0.17642 | 0.0332 |
|  | TC | -0.17553 | 0.0341 |
|  | LDL | -0.01313 | 0.8750 |

### 3.5. Regression analysis in study subjects

Univariate and multivariate linear regression analyses were conducted to determine the association of serum adiponectin with metabolic risk factors in all participants. Univariate linear analyses showed that the level of serum adiponectin was negatively correlated with BMI (β= 0.232, p <0.0001), WC (β= –0.073, p < 0.0001), SBP (β= –0.048, p < 0.0001), DBP (β= –0.0724, p < 0.0001), FBS (β= –0.008, p < 0.0006), TG (β= –0.009, p < 0.0001) and TC (β= – 0.0193, p < 0.0001), respectively. In contrast, the correlation was positive and highly significant between adiponectin and HDL (β= 0.144, p < 0.0001) (**Table 5**).

### 3.6. Adiponectin-guided drug repurposing by molecular docking

To explore Adiponectin-guided repurposable drug molecules, we performed molecular docking Adiponectin as receptors and candidate drug molecules. The 3D structures of one receptor (Adiponectin) were taken from the Protein Data Bank (PDB) using the following PDB codes:5LXG, using their corresponding UniProt IDs, Q96A54. **Table S2** displayed the top-order 30 (Out of 159) drug molecules corresponding to the receptors. The top-ranked three potential drugs (Out of 159) were considered as potential drugs because all of them exhibited significant binding affinity (BA) < –7.0 (kcal/mol) after docking. As a result, we determined that the best pharmacological molecules to block the suggested gene (Adiponectin) were Saquinavir, Candesartan and Glimepiride **Table 7**.

### 3.7. In-silico validation of candidate drug molecules by ADME/T analysis

Three (Saquinavir, Candesartan and Glimepiride) of the three compounds identified by molecular docking study satisfied at least four of Lipinski’s rule of five factors, demonstrating their drug-like qualities (**Table 8A**). The efficacy and indemnity of the suggested compounds’ ADME and toxicity (ADME/T) studies can be assessed using a variety of parameters (**Table 8B**). These three chemical compounds have a high HIA score (≥ 50%), indicating adequate absorption in the gastrointestinal (GI) tract. Since every compound blood-brain barrier (BBB} value is less than 0.3, there is a lower chance of CNS damage with none of the substances expected to penetrate the BBB. These chemicals are not harmful, according to the toxicity studies. Three substances (Saquinavir, Candesartan and Glimepiride) would act like drugs and could be taken orally based on the Drug-Likeness and ADME/T study.

## 4. Discussion

The Met-S represents a cluster of interrelated common clinical disorders, including obesity, glucose intolerance, insulin resistance, dyslipidemia, and hypertension. It is considered as the basis for the development of atherogenic cardiovascular diseases, specifically in the population of industrially developed nations who exposed to overnutrition[2,14,31]. It is alarming that the number of patients with Met-S is increasing worldwide gradually. However, the molecular basis of the clinical cluster yet to be elucidated. Adiponectin is a protein, derived from adipose tissue, which performs multiples crucial functions, such as anti-atherogenic, anti-inflammatory, lipid oxidation enhancing, insulin-sensitizing, and vasodilatory function [15,23,32–37]. Therefore, reduced levels of serum adiponectin might lead to the progression of metabolic disorders.

In this cross-sectional analysis, we found that the serum adiponectin levels ranged from 1.092 µg/mL to 10.782 µg/mL and were significantly correlated with each component of Met-S (**Figure 2**). In addition, the prevalence of multiple risk factors was higher in the group of populations with lower serum adiponectin levels (**Table 4**, **Figure 2**). This result might be a useful biomarker for the diagnosis of Met-S. [14] have reported that serum adiponectin levels are significantly lower in patients with Met-S compared to the control group. Similarly, they demonstrated that lower adiponectin concentration was associated with different components of the Met-S. The findings of our experiment also support a previous study conducted by Ahmed [38]

**Table 4.** The relation between Met-S components and serum adiponectin levels.

| Number of components | Number of observations | Serum adiponectin |  |  | SD |
| --- | --- | --- | --- | --- | --- |
|  |  | Mean | Lower | Upper |  |
| | | ($\mu\text{g/ml}$) | ($\mu\text{g/ml}$) | ($\mu\text{g/ml}$) | |
| 0 | 20 | 5.9214 | 4.9221 | 6.9206 | 2.135 |
| 1 | 44 | 5.8539 | 5.2209 | 6.4869 | 2.0821 |
| 2 | 58 | 5.2112 | 4.6148 | 5.8076 | 2.2681 |
| 3 | 66 | 4.5365 | 3.9741 | 5.099 | 2.2879 |
| 4 | 76 | 3.7281 | 3.2999 | 4.1562 | 1.8736 |
| 5 | 5 | 3.7026 | 2.5021 | 4.9031 | 0.9669 |

**Table 5.** The correlation was positive and highly significant between adiponectin and HDL (β= 0.144, p < 0.0001).

| Variables | $\beta$ | SE | Adjusted R<sup>2</sup>-value | P-value |
| --- | --- | --- | --- | --- |
| Age | -0.00436 | 0.01617 | -0.0035 | 0.7877 |
| BMI | -0.23244 | 0.04790 | 0.0776 | <.0001 |
| WC | -0.07350 | 0.01253 | 0.1109 | <.0001 |
| SBP | -0.04841 | 0.00899 | 0.0946 | <.0001 |
| DBP | -0.07243 | 0.01384 | 0.0897 | <.0001 |
| FBS | -0.00878 | 0.00254 | 0.0393 | 0.0006 |
| TG | -0.00976 | 0.00112 | 0.2193 | <.0001 |
| HDL | 0.14467 | 0.01920 | 0.1723 | <.0001 |
| TC | -0.01933 | 0.00433 | 0.0660 | <.0001 |
| LDL | -0.00657 | 0.00559 | 0.0014 | 0.2409 |

**Table 6.**
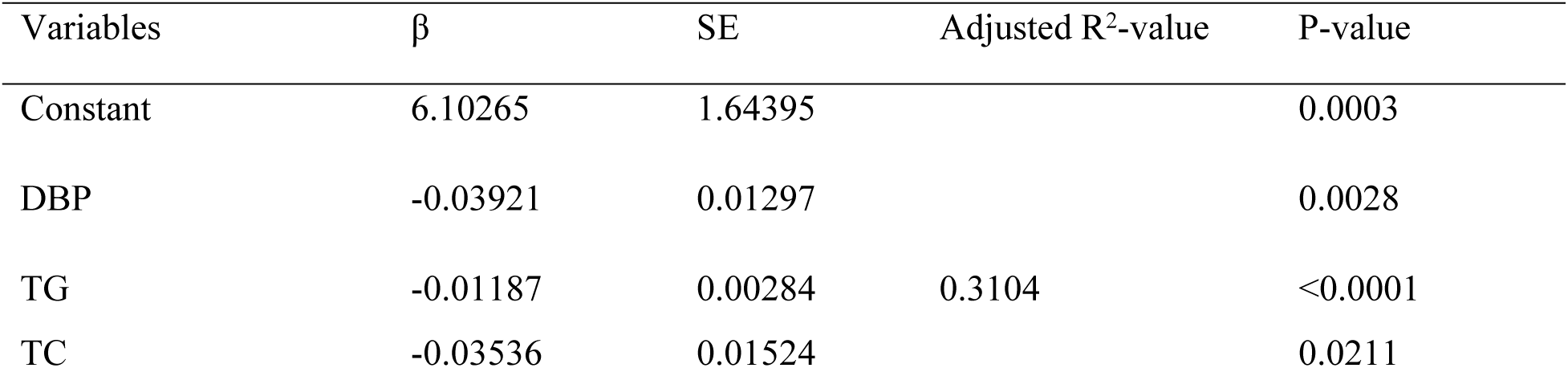
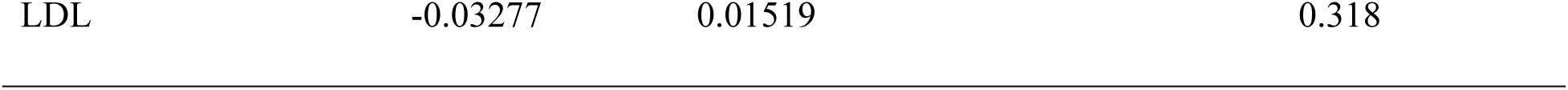
Multivariate linear regression of serum adiponectin with clinical and biochemical parameters.

**Table 7.**
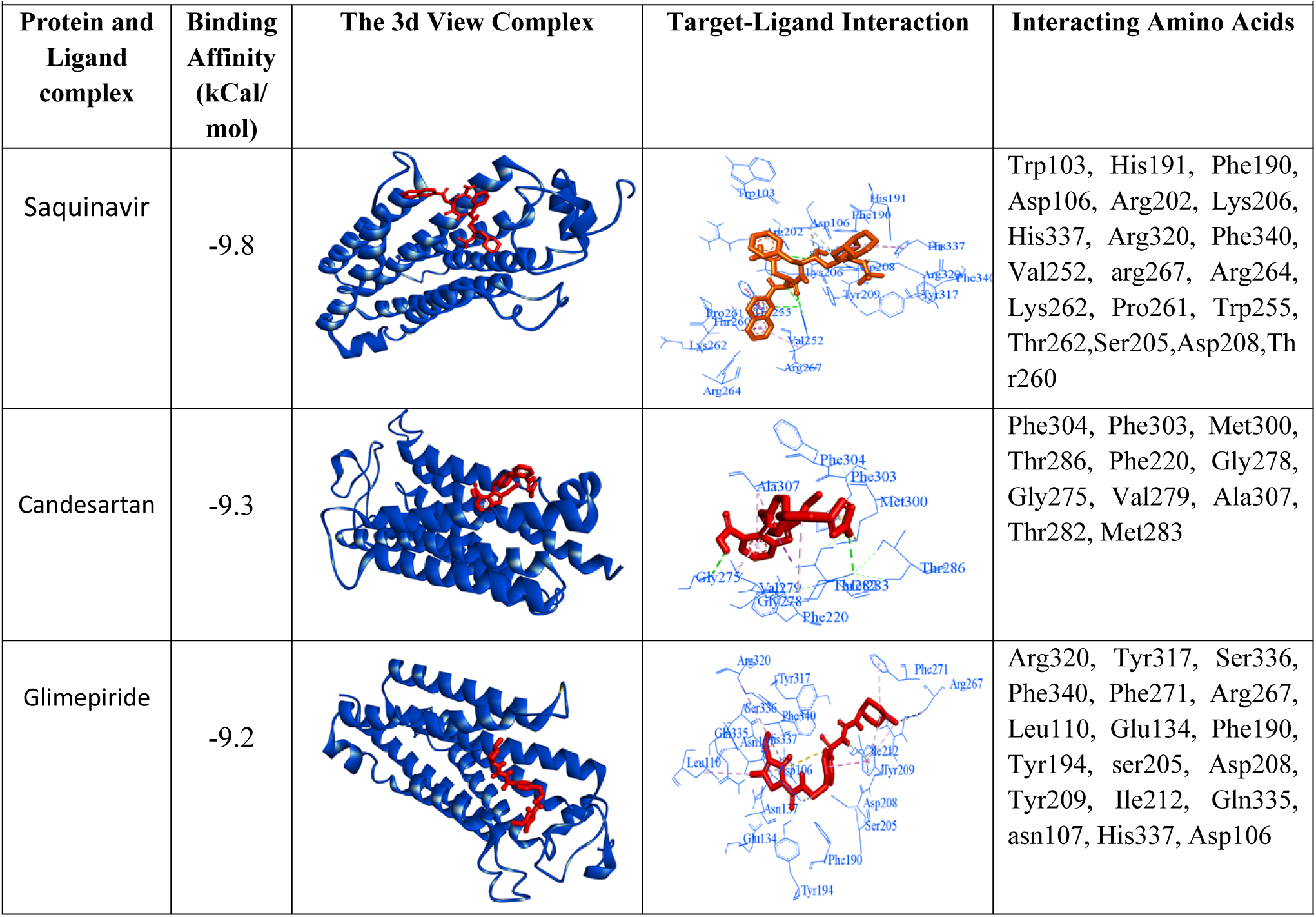
The 3-dimension view of strong binding interactions between targets and drugs.

**Table 8A.**
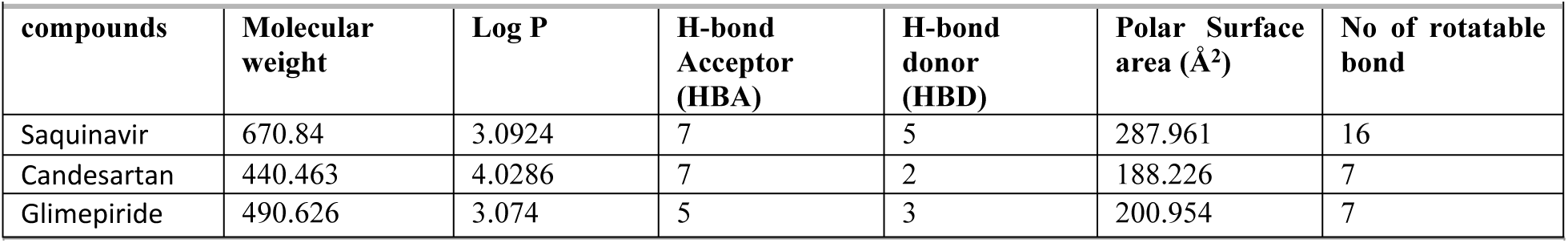
Drug likeness profile of candidate drug molecules.

**Table 8B.**
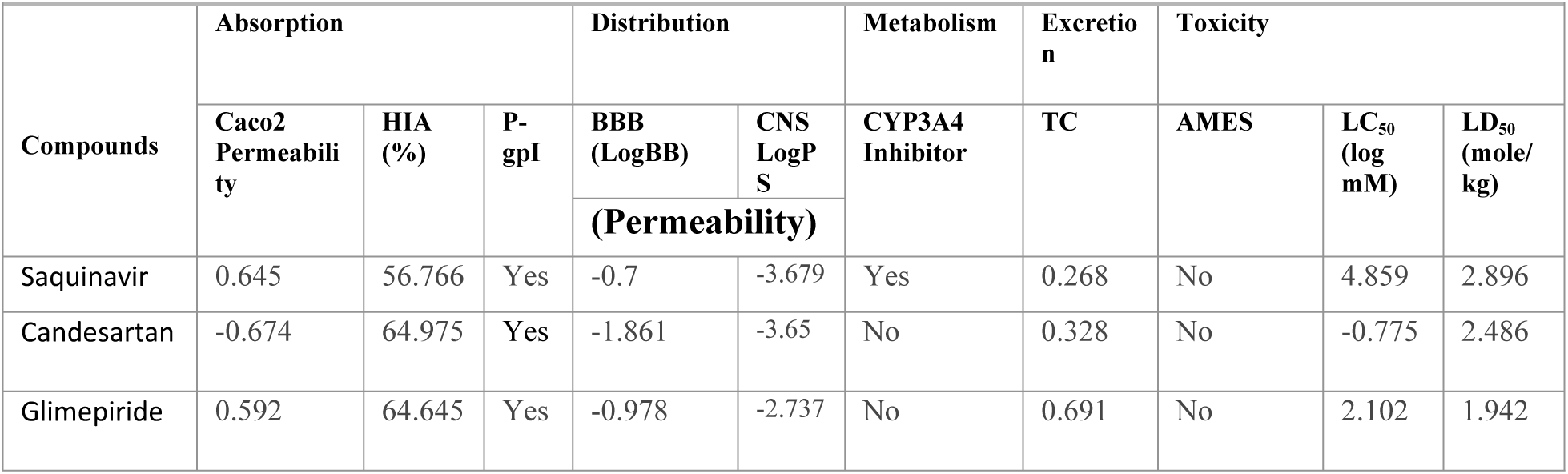
ADME and Toxicity (ADME/T) profile of top-ranked 3 drug molecules.

Our study demonstrates that there are significant differences (p < 0.0001) in the value of several parameters, including BMI, WC, SBP and DBP, FBS, TAG, and TC, between these two groups, in which the values of the above parameters are higher in participants with Met-S than those of without Met-S. Moreover, the HDL levels are decreased in Met-S patients compared to the control group (p < 0.0001). Recently, Zhuo [39] have reported lower levels of serum adiponectin in subjects with Met-S compared to individuals without Met-S. They have further concluded that lower values of adiponectin are associated with different components of the Met-S, and the adiponectin levels decrease with the increasing numbers of Met-S components, and it supports our findings.

We have identified the correlation of serum adiponectin with parameters of Met-S. The results reveal that the individuals with hypertriglyceridemia or decreased HDL-cholesterol have considerably lower serum adiponectin levels than without Met-S subjects. The outcomes of our analyses are consistent with the findings of other groups [14,38,40]. Although the specific correlation between adiponectin and lipid metabolism is unknown, it has been suggested that adiponectin may accelerate cholesterol transport in reverse order by boosting HDL assembly in the hepatic cell [41,42]. Besides, since hypoadiponectinemia is significantly associated with hypertriglyceridemia as well as reduced HDL cholesterol, this finding suggests that increased adiponectin levels may exhibit protective effects by either upregulating HDL cholesterol or downregulating triglycerides levels.

Among individuals of group 1, serum adiponectin shows significant negative correlations with WC, BMI, DBP, TG, and slightly negative correlation with SBP. However, adiponectin exhibits a significant positive correlation with serum HDL (p = 0.033), which is supported by a recent report [43]. We have not found significant correlation between serum adiponectin levels and FBS. These results contradict recent findings [14,38,43]. This confliction may be due to the formation of high molecular weight (HMW) adiponectin multimers [44]. This HMW adiponectin may be an active conformation of the adiponectin molecule and can be linked to the progression of Met-S. Details study are necessary to unveil the problem. We could not exclude the possibility for this discrepancy that most of our study subjects with Met-S were diabetic and were taking hypoglycemic drugs.

We investigated potent drugs for the treatments against Met-S and found four drugs (Saquinavir, Candesartan, and Glimepiride) displayed favorable profiles. Among the top three identified candidate drugs molecules, Saquinavir, Glimepiride and Saquinavir received support as the common candidate molecules for Met-S (diabetes, obesity etc). It should be noted here that both drug molecules are already approved by FDA by using Candesartan medication of hypertension, Glimepiride used to treat type 2 diabetes mellitus, and Saquinavir for the treatment of HIV-1, which can be found with Drug Bank (DB) accession ID DB13919, DB00222 and DB01232, respectively. According to the reference article indicate that, candesartan may be able to prevent blood pressure[45] diabetes, hypertension, heart failure[46] left ventricular hypertrophy and metabolic syndrome[47,48]. A recent study showed that glimepiride was beneficial for obese patients with T2D on both stages of insulin secretion [49]. Thus, glimepiride remains an important therapeutic option for T2D[50]. Another article shows that, Hypoxia-induced pulmonary hypertension in rats was treated with saquinavir. This study was conducted among the urban population. Since the socioeconomic status of individuals living in rural and urban areas varied greatly, the results of this study cannot be generalized for the whole Bangladeshi population. A nationwide study with a large sample size from different socioeconomic backgrounds may shed light on the function of adiponectin in the progression of Met-S. Increasing sample size and turning this cross-sectional study into cohort research with a longer follow-up period may minimize the research disparity. A change in lifestyle may aid in the management of serum adiponectin levels. Continued weight loss programs of diabetic, non-diabetic and obese individuals may increase serum adiponectin levels.

## 4. Conclusion

Based on our findings, we can conclude that plasma adiponectin levels are significantly lower in individuals with metabolic syndrome (Met-S) compared to those without Met-S. Furthermore, lower adiponectin levels are associated with specific Met-S risk factors such as hypertriglyceridemia and reduced HDL-cholesterol. These results suggest that adiponectin may serve as a potential diagnostic and prognostic marker for Met-S. Finally, adiponectin targeted top-ranked three candidate drug agents (Saquinavir, Candesartan and Glimepiride) were discovered through molecular docking, and ADME/T analysis.

## Declarations

### Credit authorship contribution statement

**Salina Shaheen Parul:** Designed the projects, performed the experiments, collected the data, analyzed the data, Writing-original draft. **Reaz Ahmmed:** Writing-original draft, Performed the experiments, Review and edited the original draft, Software and Validation. **Md. Taohid Hasan:** Writing-original draft, Performed the experiments, Review and editing the original draft. **Ariful Islam:** Review the original draft, **Md. Motiar Rahman:** Review and Editing **M. Manirujjaman:** Review and Editing. **Md. Wasim Bari:** Edited and review the original draft and **Md. Shakil Ahmed:** Review the original draft. **Md. Sohel Hasan:** Review the original draft, **Mohammad Amirul Islam:** Conceptualization, Supervision, Resources, Project Administration, Writing-review & editing.

### Ethical approval

The study was conducted in compliance with the regulations of the Ethical Review Committee (ERC) (Ref. KMC/ERC/034(A)) of Khulna Medical College (KMC) on 21 September 2016.

### Data availability statement

Data will be made available on request.

### Consent for publication

All the authors approved the final version of the manuscript for publication.

### Additional information

No additional information is available for this manuscript.

## Acknowledgements

The authors are grateful to the Department of Medicine of Khulna Medical College Hospital for collecting the patient’s sample & performing the experiments and also to the Department of Biochemistry and Molecular Biology of University of Rajshahi. This project was partially financially supported by the Institute of Biological Sciences, University of Rajshahi. It also slightly funded by the faculty of Science University of Rajshahi.

